# Evaluation of liver protective activity of *Moringa oleifera* bark extract in paracetamol induced hepatotoxicity in rats

**DOI:** 10.1101/513002

**Authors:** Rajibul Islam, Md. Jahir Alam

## Abstract

**Background:** *Moringa oleifera* has been used in folk medicine to alleviate several diseases. In the present study, ethanolic extract of *Moringa oleifera* bark has been investigated to study its potential on paracetamol induced hepatotoxicity on model rats.

**Methods:** Rats (150–200 gm) were divided into 5 groups containing 6 animals each. Acute hepatotoxicity was induced by paracetamol (600 mg/kg body weight) administered once daily for one week whereas the extract of investigated plant was given orally throughout the whole experiment at 250 and 500mg/kg body weight. Silymarin (100mg/kg body weight) was given orally as standard hepatoprotective drug. The level of hepatic injury recovery was determined by the estimation of liver enzymes like SGPT, SGOT, ALP, Bilirubin, Total protein and Albumin.

**Results:** Treatment with MO extract as well as standard hepatoprotective agent silymarin ameliorated the increased plasma levels of these hepatic enzymes and indicated the hepatoprotective potential of the extract.

**Conclusion:** The biochemical parameters provide evidence that the ethanolic extract of of *Moringa oleifera* bark has shown hepatoprotective activity.

## Introduction

Liver is the major organ which plays a key roles in processing critical biochemical and physiological phenomena including metabolism and Detoxification of endogen and exogenous compounds, such as drugs and xenobiotics, homeostasis, growth, energy and nutrient supply [1, 2] Hepatic injury could be occurred by hepatotoxic agents including drugs, alcohol and viral infections.[1,3] Liver diseases like jaundice, cirrhosis and fatty liver have been public health concern across the world. Prevalence of chronic liver disease worldwide is 18.5% and cirrhosis is 4.5 to 9.5% while 2 million people die each year [4]. In terms of medication, conventional or synthetic drugs are limited. Moreover they can have serious side effects [5]. Due to the fact, a huge number of medicinal plants have been trialed to figure out regenerative and hepatoprotective activity [3] Approximately 160 phytochemical constituents originated from 101 plants have been reported to be potentially hepatoprotective.[6] At present time, medicinal herbs have been a vital source of treatment of liver disease for instance, hepatitis, cirrhosis, and loss of appetite [7].Bangladesh, a country of great biodiversity of medicinal plants having a long history of use of traditional medicine along with phytotherapeutic potential mainly from native source. As a result, research in medicinal plants have been a huge area of discovering of promising lead compounds in Bangladesh.[8] *Moringa oleifera*, a medicinal plant of Bangladesh, belongs to the family moringaceae, is a small or middle sized tree, usually grows 10-12 m in height. The plant is indigenous and abundantly seen in India, Pakistan, Bangladesh and Afghanistan. *M. oleifera* produce drumstick-like fruits. The flowers are white and quite small. It has teardrop shaped round and small leaves. Fruits and leaves are edible which are generally eaten as green vegetable. [9][10] A good number of phytochmicals has been isolated and reported from various parts of *M. oleifera* which are 4-[(4’-O-acetylalpha-Lrhamnosyloxy) benzyl isothiocyanate, Niaziminin A, and Niaziminin B, 4-(alpha-1-rhamnopyranosyloxy)-benzylglucosinolate, quercetin-3-O-glucoside and quercetin-3-O-(6”-Malonyl-glucoside),Niazimicin (pyrrolemarumine400-O-a-L-rhamnopyranoside) and 40-hydroxyphenylethanamide (marumoside A and B), Isothiocyanate, nitrites, thiocarbamates,O-(1heptenyloxy) propyl undecanoate, O-ethyl-4-(alpha-L-rhamnosyloxy) benzyl carbamate, methyl-p-hydroxybenzoate, beta-sitosterol, 4-hydroxyl mellein, vanillin, octacosonoic acid, Methionine, cysteine, D-glucose, kaemopherol, kaempferitin and ascorbic acid, protein, D-mannose, Moringine, moringinine, spirachin, 1,3-dibenzyl urea, alpha-phellandrene, p-cymene, Deoxy-niazimicine [10,11, 12] A great number of pharmacological activity has also been reported. Aqueous and alcoholic extracts of leaves and roots of *Moringa oleifera* has been reported to have a strong *in-vitro* anti-oxidant and radical scavenging activity[12] while Methanolic and ethanolic extract of leaves against pentylenetetrazole and maximal electroshock induced convulsions, demontrated significant anti-convulsant activity [13,14] Potent anti-diabetic activity from different extracts of leaves, seed and pod, has also been reported [,15,16, 17] an investigation against seed power also revealed its Anti-asthmatic activity [19,18] Study on Ethanolic extracts of *Moringa oleifera* exhibited potent anti–tumor [19] and Anthelmintic activity [21, 22] Another study of alcoholic extracts of leaves as well as seeds has showed effectiveness on isoniazid, rifampicin, and pyrizinamide induced liver damage.[23] However, our present study was concentrated on the bark extract of *Moringa oleifera* on paracetamol induced liver damage on Sprague Dawley rats.

## MATERIALS AND METHODS

### Plant collection and Extraction

The plant *Moringa oleifera* (MO) bark was collected from Savar area and was identified and authenticated by the Dept. of Botany, Jahangirnagar University, Savar, Dhaka. The collected materials were thoroughly washed in water, cut into smaller parts and shed dried at 35° – 40° C for a week and pulverized in electric grinder to get extractable powder. Then powder was extracted in soxhlet apparatus with ethanol (96%). Then the solvent washing the constituents of powder was collected in a container, dried with a rotary evaporator under reduced pressure to get viscous substance. Finally a solid mass was obtained and preserved in a petridis in the refrigerator.

### Experimental animals

For the experiment, Sprague Dawley rats of either sex, weighing between 150-200g, were collected from the animal research lab in the Department of Pharmacy, Jahangirnagar University, Savar, Dhaka. Animals were maintained under standard environmental conditions (temperature: 27.0±1.0°, relative humidity: 55-65% and 12 h light/12 h dark cycle) and had free access to feed and water *ad libitum*. The animals were acclimatized to laboratory condition for one week prior to experiments. All protocols for animal experiment were approved by the institutional animal ethical committee.

### Toxicity studies

Toxicity studies of the extracts were carried out in Swiss Albino mice of either sex weighing between 20 and 25 g. The extract was found to be safe till 5000 mg/kg p.o. Therefore, doses were selected as 250 mg/kg and 500 mg/kg b.w. [24].

### Experimental design for the assessment of liver functions

Animal study was performed at Pharmacology Laboratory, Department of Pharmacy, Jahangirnagar University, Savar, Dhaka-1342. The rats were housed in polypropylene cages at room temperature (27±2°C). The rats were divided into five groups of 6 animals (n = 6) each [25].

**Group I:** received water (10 mL/kg p.o.) once daily for 7 days, and served as normal control

**Group II:** received water (10 mL/kg p.o.) once daily for 7 days and served as paracetamol control.

**Group III:** received standard drug silymarin (100mg/ kg p.o.). once daily for 7 days, serving as STD.

**Group IV and V:** received *Moringa oleifera* bark extract (250 and 500 mg/kg respectively) once daily for 7 days. In all groups except group-I paracetamol 600mg/kg bw p.o was administered once daily with respective treatment according to Tabassum N and Agrawal SS (2004) with slight modification with error and trial [26].

Rats were anesthetized using ketamine (500mg/kg, i.p.). After sacrifice, blood samples from each group of rats were collected and the serum was separated by centrifugation. Serum samples were subjected to liver function tests of enzymes such as glutamate pyruvate transaminase (GPT/ALT), glutamate-oxaloacetate tranaminase (GOT/AST) [27], alkaline phosphatase (ALP) [28], total bilirubin [29] and total protein by standard enzymatic colorimetric method.

### Statistical Analysis

Statistical analysis for animal experiments was carried out using One way ANOVA following Bonferroni’s post hoc test using SPSS 16.0 for windows. Data were presented as Mean ± SEM. The results obtained were compared with the vehicle treated paracetamol control group. *p* values <0.05, <0.01 and <0.001 were considered to be statistically significant, highly significant and very highly significant respectively.

## Result and Discussion

Acetaminophen (AAP) is a frequently used analgesic that causes the formation of NAPQI and hepatic damage by GSH depletion. At high doses, more NAPQI will bind covalently to cellular macromolecules [30, 31, 32, 33]. Therefore, macromolecules like enzymes will leak from the damaged tissues into the bloodstream, [34] and a study of these enzyme activities in plasma has been found to be of great importance in the assessment of liver damage [35].

The increased levels of AST and ALT indicate cellular damage and loss of functional integrity of the hepatocytes [36]. The increase in ALP in liver disease reflects the pathological alteration in biliary flow [37].

The present study demonstrated that the MO extract decrease the SGPT very highly significantly (p<0.001), SGOT level highly significantly (p<0.01) at 500 mg/kg dose and also the ALP level significantly at 250 mg/kg (p<0.01) and 500 mg/kg dose (p<0.001) (table 2 and figure 2).

**Figure 1.1:**
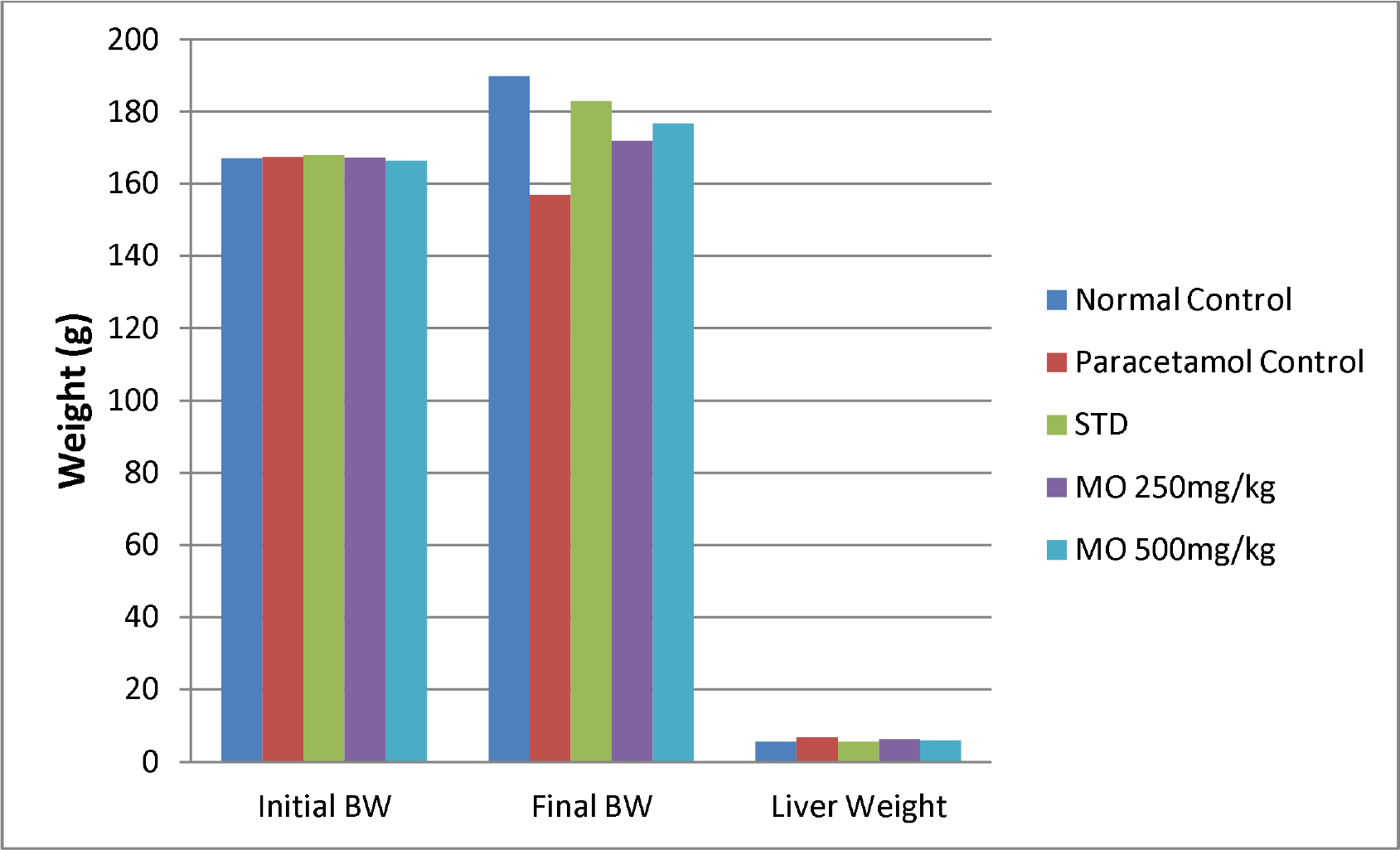
Effect of *Moringa oleifera* bark on the body weight and liver weight in paracetamol-induced hepatotoxicity in rats

**Figure 2.1:**
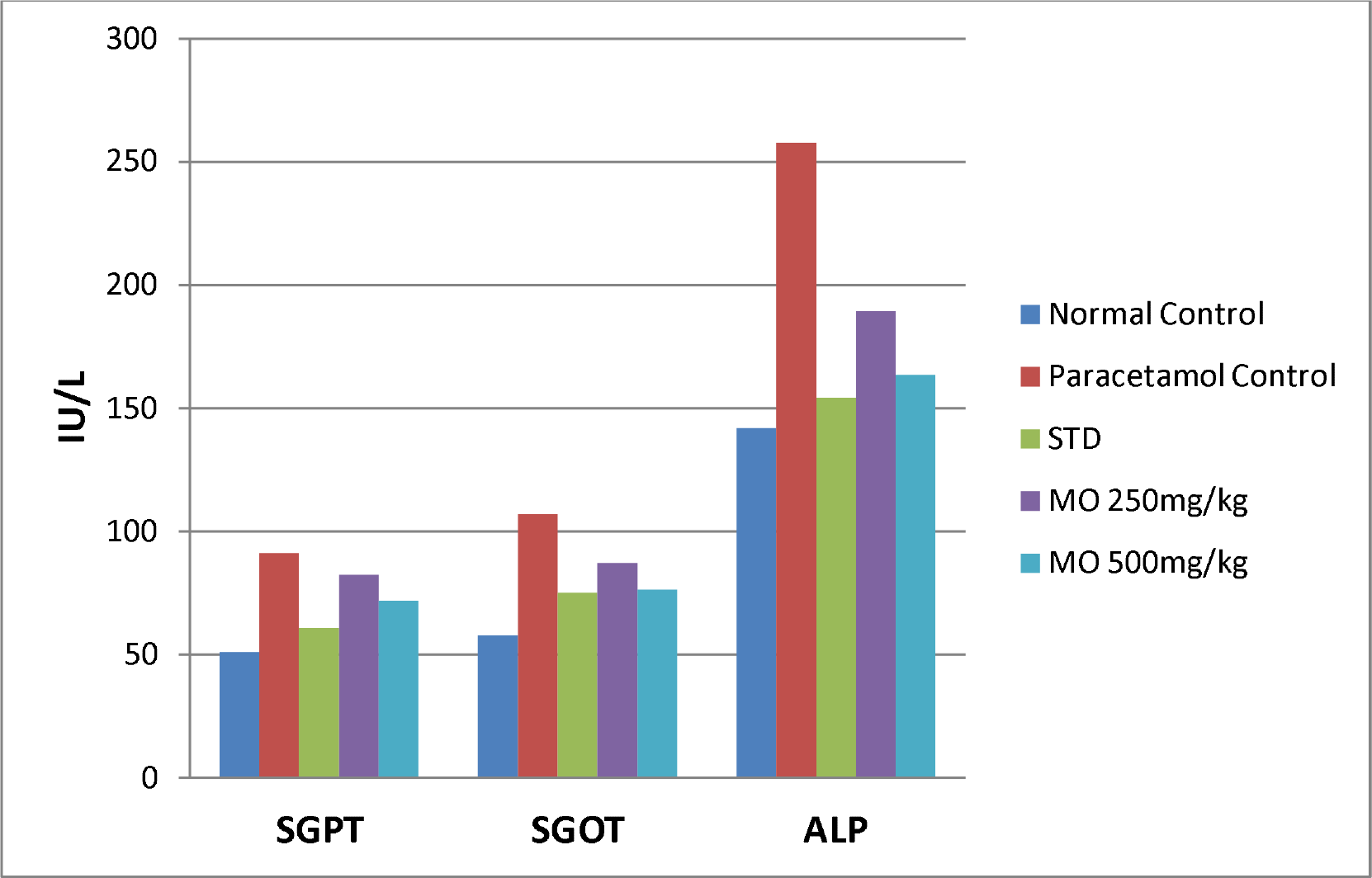
Effect of *Moringa oleifera* bark on SGPT, SGOT and ALP level in paracetamol-induced hepatotoxicity in rats.

**Figure 2.2:**
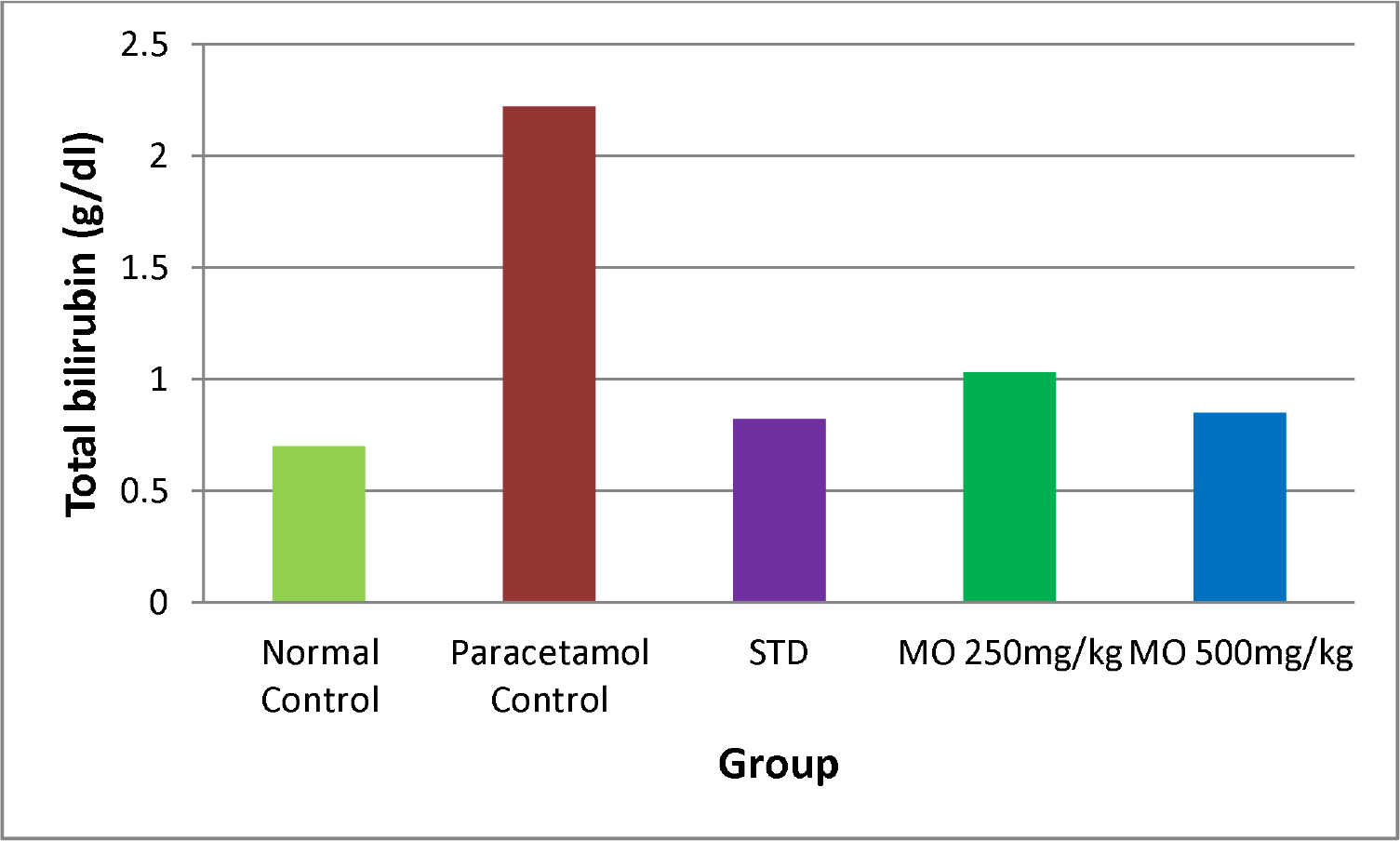
Effect of *Moringa oleifera* bark on serum bilirubin level in paracetamol-induced hepatotoxicity in rats.

**Table 1:**
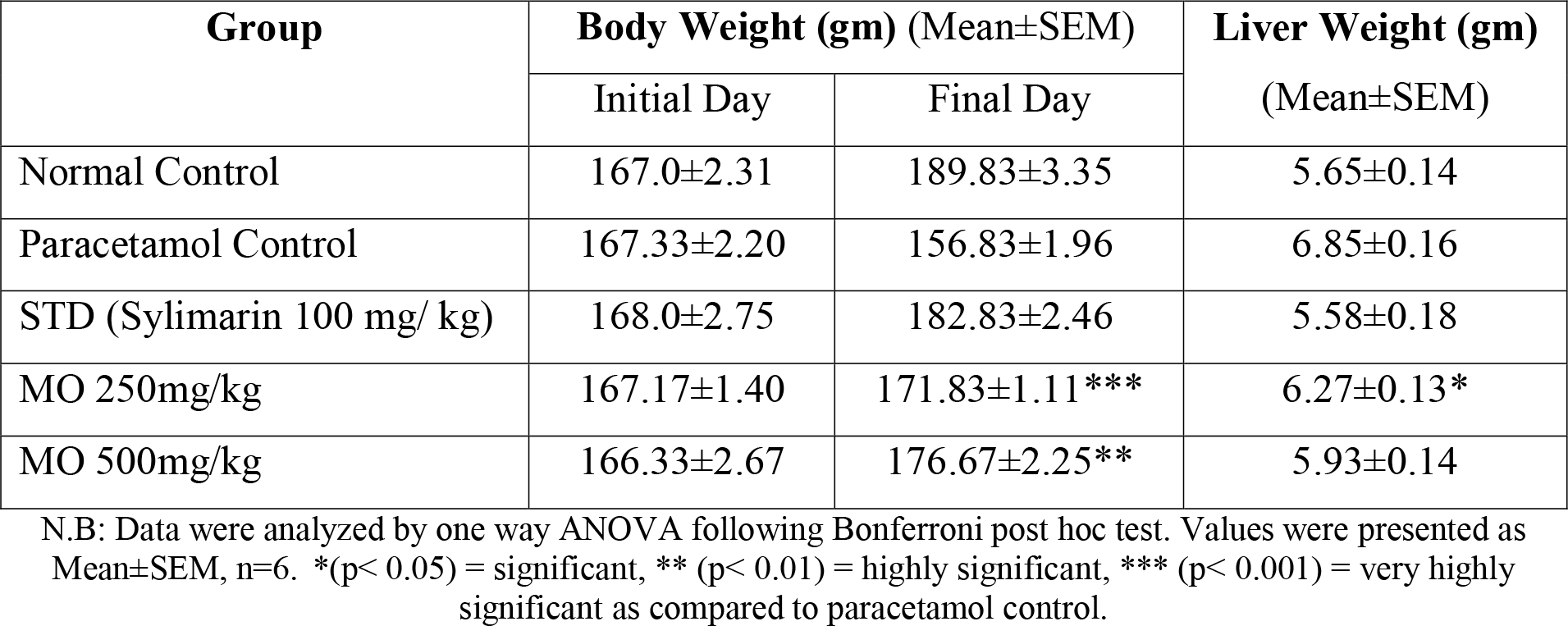
Effect of *Moringa oleifera* bark on the body weight and liver weight in paracetamol-induced hepatotoxicity in rats.

**Table 2:**
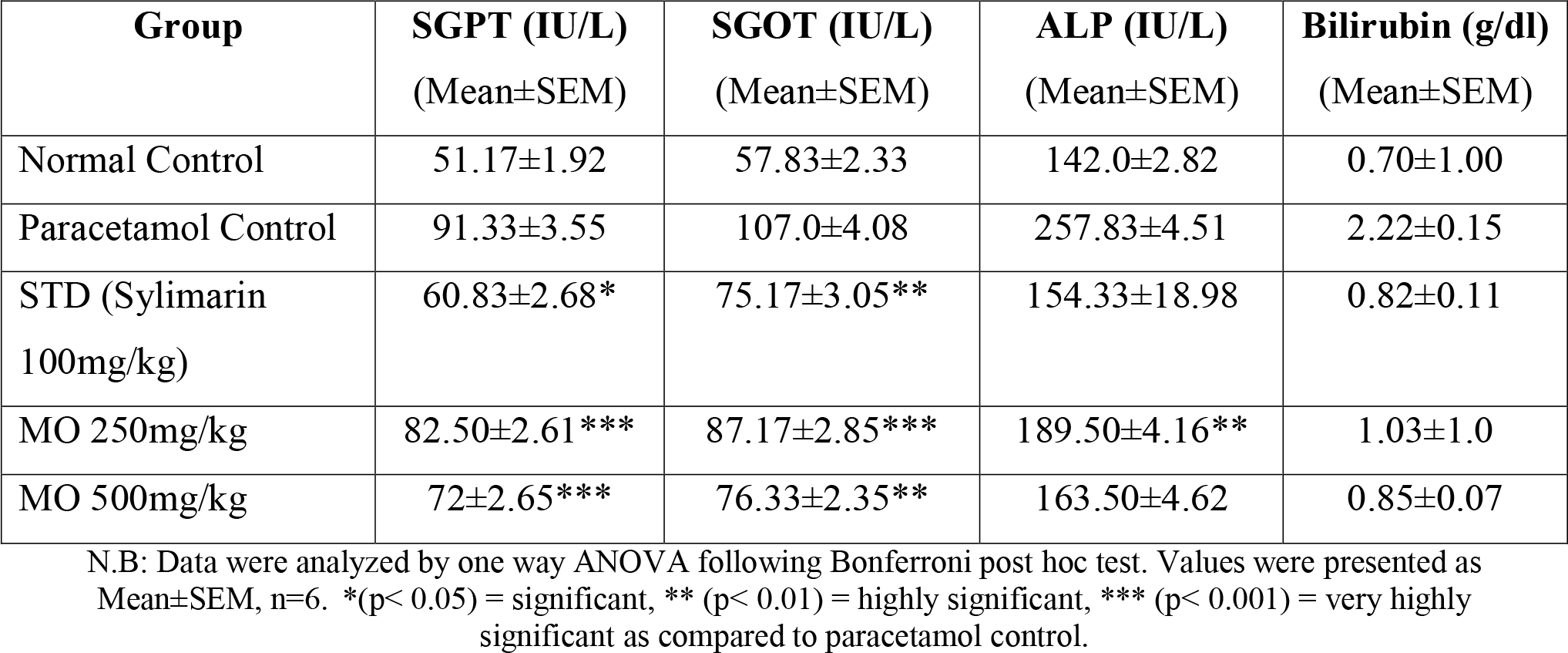
Effect of *Moringa oleifera* bark on serum GPT, GOT, ALP and bilirubin level in paracetamol-induced hepatotoxicity in rats.

| Group | SGPT (IU/L)<br>(Mean±SEM) | SGOT (IU/L)<br>(Mean±SEM) | ALP (IU/L)<br>(Mean±SEM) | Bilirubin (g/dl)<br>(Mean±SEM) |
| --- | --- | --- | --- | --- |
| Normal Control | 51.17±1.92 | 57.83±2.33 | 142.0±2.82 | 0.70±1.00 |
| Paracetamol Control | 91.33±3.55 | 107.0±4.08 | 257.83±4.51 | 2.22±0.15 |
| STD (Sylmarin<br>100mg/kg) | 60.83±2.68* | 75.17±3.05** | 154.33±18.98 | 0.82±0.11 |
| MO 250mg/kg | 82.50±2.61*** | 87.17±2.85*** | 189.50±4.16** | 1.03±1.0 |
| MO 500mg/kg | 72±2.65*** | 76.33±2.35** | 163.50±4.62 | 0.85±0.07 |

N.B: Data were analyzed by one way ANOVA following Bonferroni post hoc test. Values were presented as Mean±SEM, n=6. \*(p< 0.05) = significant, \*\* (p< 0.01) = highly significant, \*\*\* (p< 0.001) = very highly significant as compared to paracetamol control.

Reduction of the enhanced level of serum SGPT, SGOT, ALP and total bilirubin by MO extract seemed to offer protection and maintain the functional integrity of hepatic cells.

An abnormal increase in the levels of bilirubin in plasma indicates hepatobiliary disease and severe disturbance of hepatocellular function [38]. Prior oral administration of *Moringa oleifera* extract exhibited significant protection against AAP-induced hepatotoxicity. It decreased the levels of bilirubin although it was insignificant at both the doses which is an indication of protection against hepatic damage caused by AAP (figure 2.4).

Plasma proteins are mainly produced by the liver, the principle exception being immunoglobulins. Severe liver damage decreases the production of various proteins resulting in reduced serum levels of total protein, albumin, and/ or globulin [39, 40]. Decreased protein production may render other abnormal test values. e.g. depletion of coagulation factors (all are globulins) may result in prolonged prothrombin or activated partial thromboplastin times [41].

The results indicate that protein level was slightly increased at 250 mg/kg and 500mg/kg dose which was insignificant. The serum albumin level was also slightly and insignificantly increased at both the doses although the globulin level was decreased (table 3 and figure 3).

**Table 3:** Effect of *Moringa oleifera* bark on serum total protein, albumin and globulin level in paracetamol-induced hepatotoxicity in rats.

| Group | T. Protein (g/dl)<br>(Mean±SEM) | Albumin (g/dl)<br>(Mean±SEM) | Globulin (g/dl)<br>(Mean±SEM) |
| --- | --- | --- | --- |
| Normal Control | 5.32±0.13 | 2.78±0.13 | 2.53±0.11 |
| Paracetamol Control | 4.72±0.15† | 1.82±0.11 | 2.90±0.19 |
| STD (Sylmarin 100mg/ kg) | 6.15±0.21** | 3.27±0.11* | 2.88±0.20 |
| MO 250mg/kg | 4.82±0.09 | 2.40±0.08 | 2.42±0.11 |
| MO 500mg/kg | 5.33±0.11 | 3.10±0.10 | 2.23±0.17 |

N.B: Data were analyzed by one way ANOVA following Bonferroni post hoc test. Values were presented as Mean±SEM, n=6. \*(p< 0.05) = significant, \*\* (p< 0.01) = highly significant, \*\*\* (p< 0.001) = very highly significant as compared to paracetamol control.

**Figure 3.**
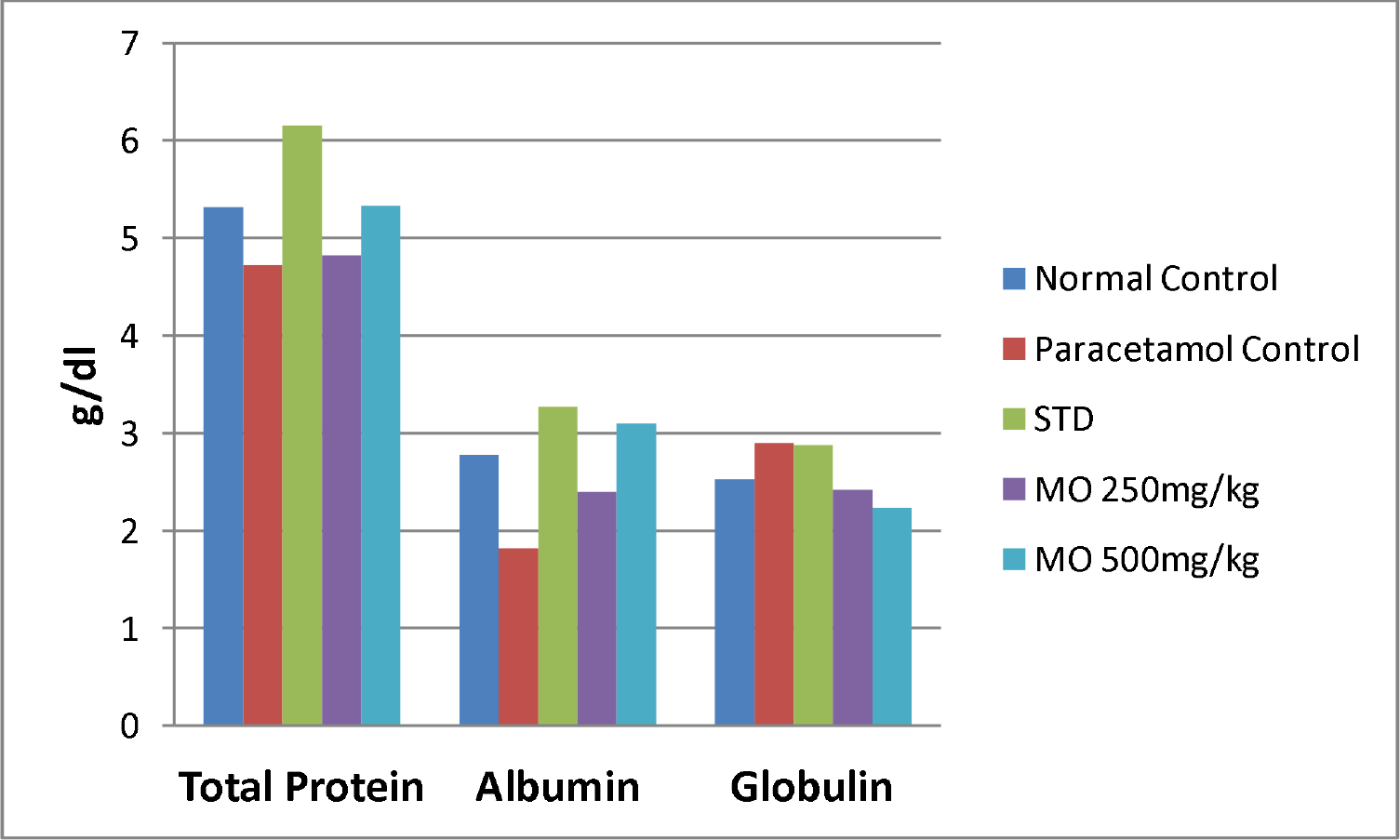
Effect of *Moringa oleifera* bark on serum total protein, albumin and globulin level in paracetamol-induced hepatotoxicity in rats.

Through this study also the consumption of MO for 7 days was found to reduce the body weight as well as the liver weight of rats (table 1 and figure 1).

## Conclusion

The above study showed that, the treatment with *Moringa oleifera* bark extract was able to protect the changes induced by AAP. On the basis of the above results it can be concluded that *Moringa oleifera* has significant hepatoprotective value against paracetamol induced liver injury. Further studies are recommended to determine the possible mechanism(s) involved.

## ACKNOWLEDGEMENTS

The author wishes to thank Pharmacology Laboratory, Jahangirnagar University, Savar, Dhaka for contribution to this research.

## References

1. Mahmood DN, Mamat SS, Kamisan HF, Yahya F, Kamarolzaman FFM, Nasir N, Mohtarrudin N, Tohid, and Zakaria AZ. Amelioration of Paracetamol-Induced Hepatotoxicity in Rat by the Administration of Methanol Extract of Muntingia calabura L. Leaves. BioMed Research International. 2014, 1–10.

2. Saleem HT, El-Maali AN, Hassan HM, Mohamed AN, Mostafa MAN, Kahaar AE, and Tammam SA. Comparative Protective Effects of N-Acetylcysteine, N-Acetyl Methionine, and N-Acetyl Glucosamine against Paracetamol and Phenacetin Therapeutic Doses–Induced Hepatotoxicity in Rats. International Journal of Hepatology. 2018, 1–8.

3. Saleem M, Asif A, Akhtar MF, SaleemA. Hepatoprotective potential and chemical characterization of Artocarpus lakoocha fruit extract. Bangladesh J Pharmacol. 2018; 13: 90–97.

4. 10th Paris Hepatology Conference. Chronic Liver Diseases: A Huge and Neglected Public Health Burden Needing Urgent Actions; 2017.

5. Rao GM, Rao CV, Pushpangadan P, Shirwaikar A. Hepatoprotective effects of rubiadin, a major constituent of Rubia cordifolia Linn. J Ethnopharmacol. 2006;103(3):484–490.

6. Chatterjee TK. Medicinal plants with hepatoprotective properties, in Herbal Opinions, third ed. Books & Allied (P) Ltd., Calcutta. 2000;135.

7. Devaraj S, Ismail S, Ramanathan S, and Yam FM. Investigation of Antioxidant and Hepatoprotective Activity of Standardized Curcuma xanthorrhiza Rhizome in Carbon Tetrachloride-Induced Hepatic Damaged Rats. Scientific World Journal 2014, 1–8.

8. Islam R., Alam MJ, Khan SA, Douti NHK. Investigation of hepatoprotective properties of the ethanolic extract of Careya arborea Roxb bark in paracetamol induced hepatotoxicity in rats. Journal of Pharmaceutical research International. 2018, 22(4) : 1–9

9. Singh A and Navneet. Ethnomedicinal, Pharmacological and Antimicrobial Aspects of Moringa oleifera Lam.: A review. The Journal of Phytopharmacology 2018; 7(1): 45–50.

10. Paikra KB, Kumar H, Dhongade J*, Gidwani B. Phytochemistry and Pharmacology of Moringa oleifera Lam. Journal of Pharmacopuncture 2017;20[3]:194–200.

11. Foidl N, Makkar HPS, Becker K. The potential use of Moringa oleifera for agriculture and industrial uses. Managua, Nicaragua. 2001;1–20.

12. Sharma VR. Paliwal R, Sharma S. Phytochemical analysis and evaluation of antioxidant activities of hydroethanolic extract of Moringa oleifera Lam. J Pharm Res. 2011;4(2):554–7.

13. Amrutia J, Lala M, Srinivasa, Moses RS. Anticonvulsant activity of Moringa oleifera leaf. International Research Journal of Pharmacy. 2011;2(7):160–2.

14. Joy AE, Kunhikatta SB, Manikkoth S. Anti-convulsant activity of ethanolic extract of Moringa concanensis leaves in Swiss albino mice. Arch Med Health Sci. 2013;1(1):6–9.

15. Ndong M, Uehara M, Katsumata S, Suzuki K. Effects of oral administration of Moringa oleifera Lam on glucose tolerance in gotokakizaki and wistar rats. J of Clin Biochem and Nutri. 2007;40:229–33.

16. Gupta R, Mathur M, Bajaj VK, Katariya P, Yadav S, Kamal R, et al. Evaluation of antidiabetic and antioxidant activity of Moringa oleifera in experimental diabetes. J Diabetes. 2012;4(2):164–71.

17. Al-Malki AL, El Rabey HA. The antidiabetic effect of low doses of Moringa oleifera Lam. seeds on streptozotocin induced diabetes and diabetic nephropathy in male rats. Biomed Res Int. 2015;2015:DOI: 10.1155/2015/381040.

18. Agrawal B, Mehta A. Antiasthmatic activity of Moringa oleifera Lam: a clinical study. Indian J Pharmacol. 2008;40(1):28–31.

19. Mehta A, Agrawal B. Investigation into the mechanism of action Moringa oleifera for its anti-asthmatic activity. Orient Pharm Exp Med. 2008;8(1):24–31.

20. Nadkarni KM. Indian materia medica. Mumbai: Popular Prakashan; 1994. 1319 p.

21. Tayo GM, Poné JW, Komtangi MC, Yondo J, Ngangout AM, Mbida M. Anthelminthic activity of Moringa oleifera leaf extracts evaluated In vitro on four developmental stages of haemonchus contortus from goats. American Journal of Plant Sciences. 2014;5(11):1702–10.

22. Rastogi T, Bhutda V, Moon K, Aswar PB, Khadabad SS. Comparative studies on anthelmintic activity of Moringa oleifera and Vitex Negundo. Asian Journal of Research in Chemistry. 2009;2(2):181–2.

23. Mishra G, Singh P, Verma R, Kumar S, Srivastav S, Jha KK, et al. Traditional uses, phytochemistry and pharmacological properties of Moringa oleifera plant: an overview. Scholars Research Library. 2011;3(2):141–64.

24. Ji Su Kim, Jung Bong Ju, Chang Won Choi and Sei Chang Kim. Hypoglycemic and antihyperlipidemic effect of four Korean Medicinal plants in alloxan induced diabetes rats. American Journal of Biochemistry & Biotechnology;2006, Vol.2 Issue 4, p154–160.

25. Bhattacharya D, Pandit S, Mukherjee R, Das N, Sur TK : Hepatoprotective effect of Himoliv® polyherbal formulation in rats. Indian J Physiol Pharmacol 2003, 47: 435–440.

26. Tabassum N and Agrawal SS,. Hepatoprotective activity of eclipta alba hassk. Against Paracetamol induced hepatocellular damage in mice. Experimental Medicine; 2004, Vol. 11, No. 4.

27. Lowry OH, Rosebrough NJ, Farr AL, Randall RJ. J Biol Chem 1951;193:265.

28. Kalyani M, Rathi M.A, Thirumoorthi L, Meenakshi P, Guru kumar. D, Sunitha M., Gopalakrishnan. V.K. Asteracantha longifolia Inhibits Perchloroethylene – Induced Hepatic Damage in Rat. Journal of Pharmacy Research,2010, 3(7),1535–1537.

29. Yoshiyuki, K., Michinori, K., Todato, T., Shigeru, A., Hiromichi, O. Studies on Scutelariae radix. Effects on lipid peroxidation of rat liver. Chemical and Pharmaceutical Bulletin 1981, 29, 610–2617.

30. Black M: Acetaminophen hepatotoxicity. Gastroenterology 1980; 78:382–392.

31. Pacifici GM, Back DJ, Orme ML: Sulphation and glucuronidation of paracetamol in human liver: assay conditions. Biochem Pharmacol 1988;37:4405–4407.

32. Jollow DJ, Mitchell JR, Potter WZ, Davis DC, Gillette JR, Brodie BB: Acetaminophen-induced hepatic necrosis: II. Role of covalent binding in vivo. J Pharmacol Exp Ther 1973;187:195–202.

33. Potter DW, Hinson JA: Reactions of N-acetyl-p-benzoquinoneimine with reduced glutathione, acetaminophen and NADPH. Mol Pharmacol 1986;30:33–41.

34. Hearse DJ: Cellular damage during myocardial ischaemia: metabolic changes leading to enzyme leakage. In: Enzymes in Cardiology (Hearse DJ, De Leiris J, Loisance D, eds.), John Wiley and Sons Ltd., New York, 1979, pp. 1–21.

35. Plaa GL, Zimmerman HJ: Evaluation of hepatotoxicity: physiological and biochemical measures of hepatic function. In: ComprehensiveToxicology, Vol. 9 (McCuskey RS, Earnest DL, eds.), Cambridge University Press, Cambridge, UK, 1997, pp. 97–109.

36. Drotman RB, Lowhorn GT: Serum enzymes as indicators of chemical induced liver damage. Drug Chem Toxicol 1978;1:163–171.

37. Plaa GL, Hewitt WR: Detection and evaluation of chemically in-duced liver injury. In: Principles and Methods of Toxicology, 2nd ed. (Wallace Hayes A, ed.), Raven Press, New York, 1989, pp. 399–428.

38. Martin P, Friedman LS: Assessment of liver function and diagnostic studies. In: Handbook of Liver Disease (Friedman LS, Keeffe EB, eds.), Churchill Livingstone, Philadelphia, 1998, pp.1–14.

39. Alper, C.A. 1974. Plasma protein measurement as a diagnostic aid. N. Eng. J. Med. 291: 287–290.

40. Killingsworth, L.M. (1981). “The Role of High resolution Electrophoresis in the Clinical Evaluation of Protein Status.” Freehold, New Jersey: Worthington Diagnostics.

41. Badylak, S.F., and Van Vleet J.F., Alteration of prothrombin time and activated partial thromboplastin time in dogs with hepatic disease. Am. J. Vet. Res. 1981, 42: 2053–2056.

